# Positive but variable effects of crop diversification on biodiversity and ecosystem services

**DOI:** 10.1101/2020.09.30.320309

**Authors:** D. Beillouin, T. Ben-Ari, E. Malézieux, V. Seufert, D. Makowski

## Abstract

Ecological theory suggests that biodiversity has a positive and stabilizing effect on the delivery of ecosystem services. Yet, the impacts of increasing the diversity of cultivated crop species or varieties in agroecosystems are still under scrutiny. The empirical evidence available is scattered in scope, agronomic and geographical contexts and impacts on ecosystem services may depend on the type of diversification strategy used. To robustly assess the effects of crop diversification in agroecosystems, we compiled the results of 95 meta-analyses integrating 5,156 experiments conducted over 84 experimental years and representing more than 54,500 paired observations on 120 crop species in 85 countries. Overall, our synthesis of experimental data from across the globe shows that crop diversification enhances not only crop production (median effect +14%), but also the associated biodiversity (+24%, i.e. the biodiversity of non-cultivated plants and animals), and several supporting and regulating ecosystem services including water quality (+51%), pest and disease control (+63%) and soil quality (+11%). However, there was substantial variability in the results for each individual ecosystem service between different diversification strategies like agroforestry, intercropping, cover-crops, crop rotation or variety mixtures. Agroforestry is particularly effective in delivering multiple ecosystem services, i.e. water use and quality, pest and diseases regulation, associated biodiversity, long-term soil productivity and quality. Variety mixtures, instead, provide the lowest benefits, while the other strategies show intermediate results. Our results highlight that while increasing the diversity of cultivated crop species or varieties in agroecosystems represents a very promising strategy for more sustainable land management, contributing to enhanced yields, enhanced biodiversity and ecosystem services, some crop diversification strategies are more effective than others in supporting key ecosystem services.

## Introduction

Driven by productivity-oriented policies, agricultural practices throughout the 20^th^ century have converged towards a strong intensification of agroecosystems^1^, which was associated with a large decrease in the number of cultivated plant species^2,3^. These intensive agricultural practices have not only resulted in a homogenization of agricultural systems, they have also resulted in agriculture becoming the major contributor of the crossing of several planetary boundaries^4^. A variety of different already existing agricultural systems have been proposed as more sustainable alternatives to current intensive systems, to ensure adequate food availability while minimizing the environmental impacts of food production^5^, including, for example, organic agriculture, agroecological farming, ecological or sustainable intensification^3,6,7^. Most of these alternative systems share the implicit or explicit goal of increasing agrobiodiversity, i.e. the diversity of cultivated crop species on-farm and in agricultural landscapes.

Ecological theory suggests that more diverse systems can provide a wider range of different ecosystem services in higher quantity and at increased stability over time compared to less diverse systems^8–10^. The mechanisms behind the proposed relationship between biodiversity, ecosystem function and the delivery of ecosystem services are manifold, complex and debated but have been summarized in four (not necessarily mutually exclusive) hypotheses: systems with higher biodiversity i) offer more different niches and thus provide more different functions (*species complementarity)*; ii) have built-in redundancy (i.e. there are more species than there are functions, *species redundancy); iii*) increase resilience, due to their redundancy, over time in the face of environmental fluctuations (*diversity-stability)*; iv) provide extremely variable relationships between a system’s biodiversity and function depending on the type and strength of species interactions (*idiosyncrasy*) ^9,11,12^ In agroecosystems, these mechanisms are further complicated, since many ecosystem functions are substituted by human labour or by agricultural inputs^8^.

The growing popularity and promise of crop diversification as a tool for more sustainable land management is evidenced in a fastly growing empirical literature (*Fig. 1a*), as well as numerous qualitative^13–19^ and quantitative^20^ synthesis. But when synthesizing this vast body of literature to provide guidance to decision-makers about the promises and shortfalls of these practices, it is crucial to (1) start with a clear definition and conceptualization of what agricultural diversification means, (2) provide a systematic and global synthesis of the current evidence base, while (3) addressing and exploring bias and heterogeneity in the data analysed, as well as (4) exploring the quality and robustness of the results.

**Figure 1.**
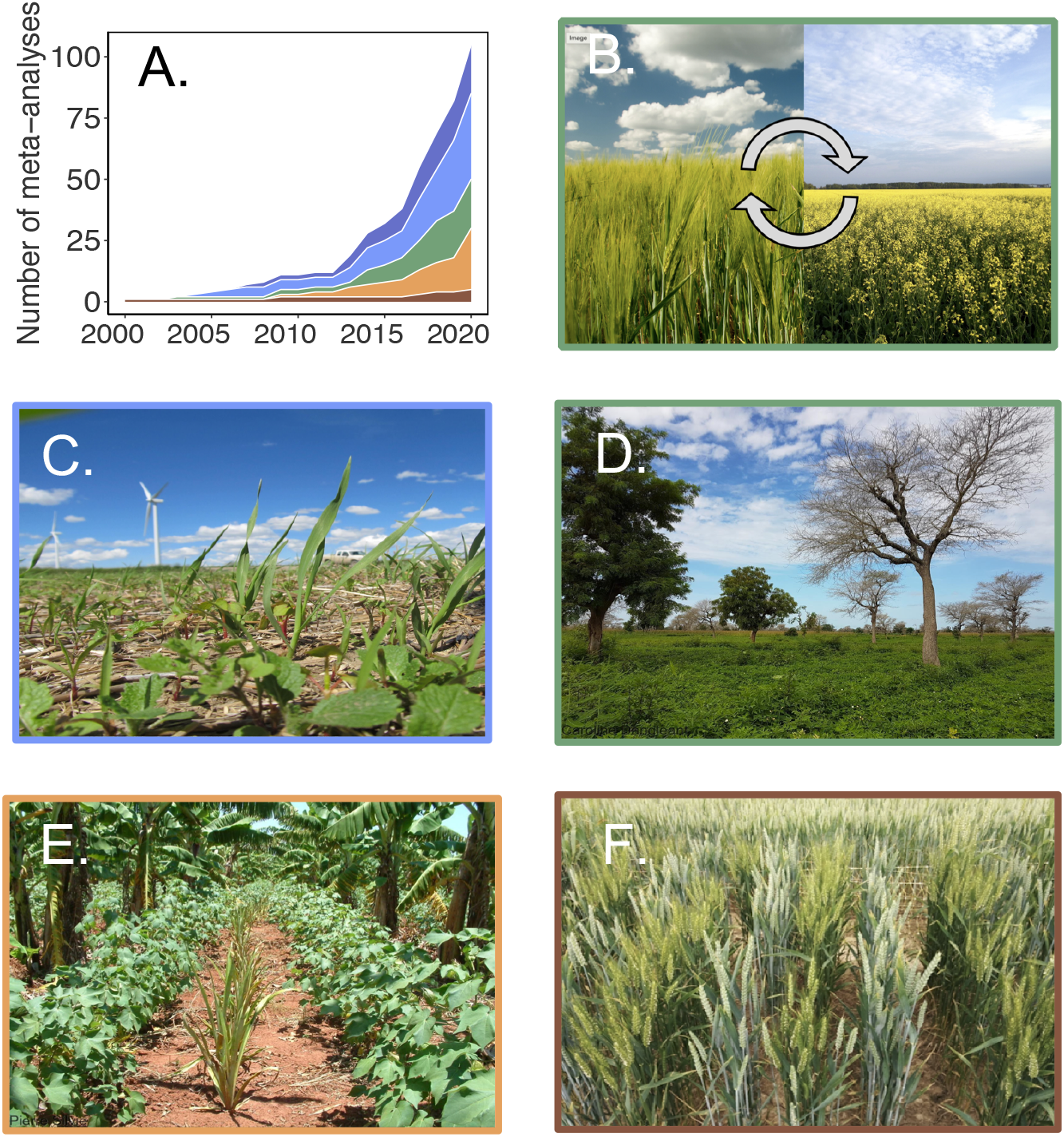
The five crop diversification strategies included in the analysis, the temporal evolution of the number of meta-analyses studying this practice (A), as well as illustrative examples of each practice (B-F). The colors of the border of each photo correspond to the color of the crop diversification strategies in panel A. The crop diversification strategies are: (B) crop rotation (example: wheat and canola rotation), (C) cover-crops (here, a mix of oats, buckwheat, turnip, radish, mustard, millet, soybean, and hairy vetch in wheat residues, USA), (D) agroforestry (here, peanuts and acacia, Senegal), (E) intercropping (here, organic cotton cultivation combined with pineapple and banana crops, Paraguay), and (F) cultivar mixture (here, wheat mixtures, France). Note that one meta-analysis could study one or multiple crop diversification strategies. For a detailed definition of each crop diversification practice see the Supplementary Materials. Credits of photo are in supplementary.

Our objective is hence to provide a rigorous global synthesis of the effects of various crop diversification practices at field scale on biodiversity and ecosystem services. We define crop diversification as any type of agricultural diversification practice that involves increasing the diversity of cultivated annual or perennial crops (including non-harvested crops like cover crops or catch crops) at genetic or species level. To allow for additional insights into the impacts of different types of crop diversification practices we group crop diversification strategies into 5 categories: *crop rotations* (i.e. temporal succession of a set of selected crops grown on a field each season or each year), *cover crops* (i.e. plant sown in addition to the main crop for agronomic or environmental purposes), *agroforestry* (i.e. inclusion of at least one woody perennial species), *intercropping* (i.e. the simultaneous cultivation in the same field of two or more different crop species for all or part of their growth cycle) and *variety mixtures* (i.e. the simultaneous cultivation in the same field of multiple varieties of the same crop species; *Fig. 1*; for a more detailed definition of the crop diversification practices see Supplementary Materials). To provide a synthesis of the empirical evidence we compile an exhaustive database of published meta-analyses on the subject and perform a meta-analysis of published meta-analyses, i.e. a second-order meta-analysis. This approach has been scarcely applied in agronomy so far, but is already common practice in the medical sciences^21^. We also focused on assessing the quality of selected meta-analyses and their redundancy (i.e., how many common primary studies are shared between distinct meta-analyses). The overall summary effect sizes computed in our analysis are weighted according to these criteria (i.e., we perform a weighted second-order meta-analysis, see methods).

## Results and Discussion

We identified 95 meta-analyses synthesizing the effects of increasing crop diversity in agrosystems at the field scale. Our dataset includes the results of 5,156 primary studies equivalent to 54,554 experiments published since 1936, spanning more than 120 crops and 85 countries in Africa, America, Asia, Europe, and Oceania. Different crop diversification practices vary in the degree they have been studied through experimental trials and ensuing meta-analyses: cover-crops (35 meta-analyses; 23,000 experiments), crop rotation (20 meta-analyses; 3400 experiments), intercropping (20 meta-analyses; 13,000 experiments), agroforestry (19 meta-analyses, 6,900 experiments), variety mixtures (5 meta-analyses; 7,800 experiments), (*Fig 1a*). These experiments and meta-analyses examined a broad suite of different agronomic and environmental outcomes, which we summarize into the following biodiversity and ecosystem services categories (for further details about how each category was defined see Supplementary Materials): agricultural production (40 meta-analyses; 23,000 experiments), soil quality (32 meta-analyses; 13,000 experiments), associated biodiversity (14 meta-analyses; 6,500 experiments), pest and disease control (12 meta-analyses; 3,100 experiments), water quality (11 meta-analyses; 1,450 experiments), water regulation (7 meta-analyses; 1,400 experiments), greenhouse gas (GHG) emissions (7 meta-analyses; 1,300 experiments), input use efficiency (5 meta-analyses; 2,000 experiments), product quality (4 meta-analyses; 1,800 experiments), profitability (3 meta-analyses; 100 experiments), yield stability (1 meta-analysis; 390 experiments).

### Crop diversification improves biodiversity and ecosystem service delivery

Considered together, the various crop diversification strategies examined improved associated biodiversity (median effect estimate equal to +24%; 95% confidence interval CI: 15%, 33%), as well as the majority of ecosystem services analyzed (*Fig. 2*), for example resulting in improved water quality (+51%; 3%, 123%), more efficient pest and disease control (+63%; CI: 31%, 103%), better soil quality (+11%; CI: 7%, 16%) and higher production levels (+14%; CI: 8%, 20%). Our results are thus in line with the positive effects found for highly diverse plant communities in natural ecosystems^22–25^ and a previous second order-meta-analysis for diversified agricultural systems^20^.

**Figure 2.**
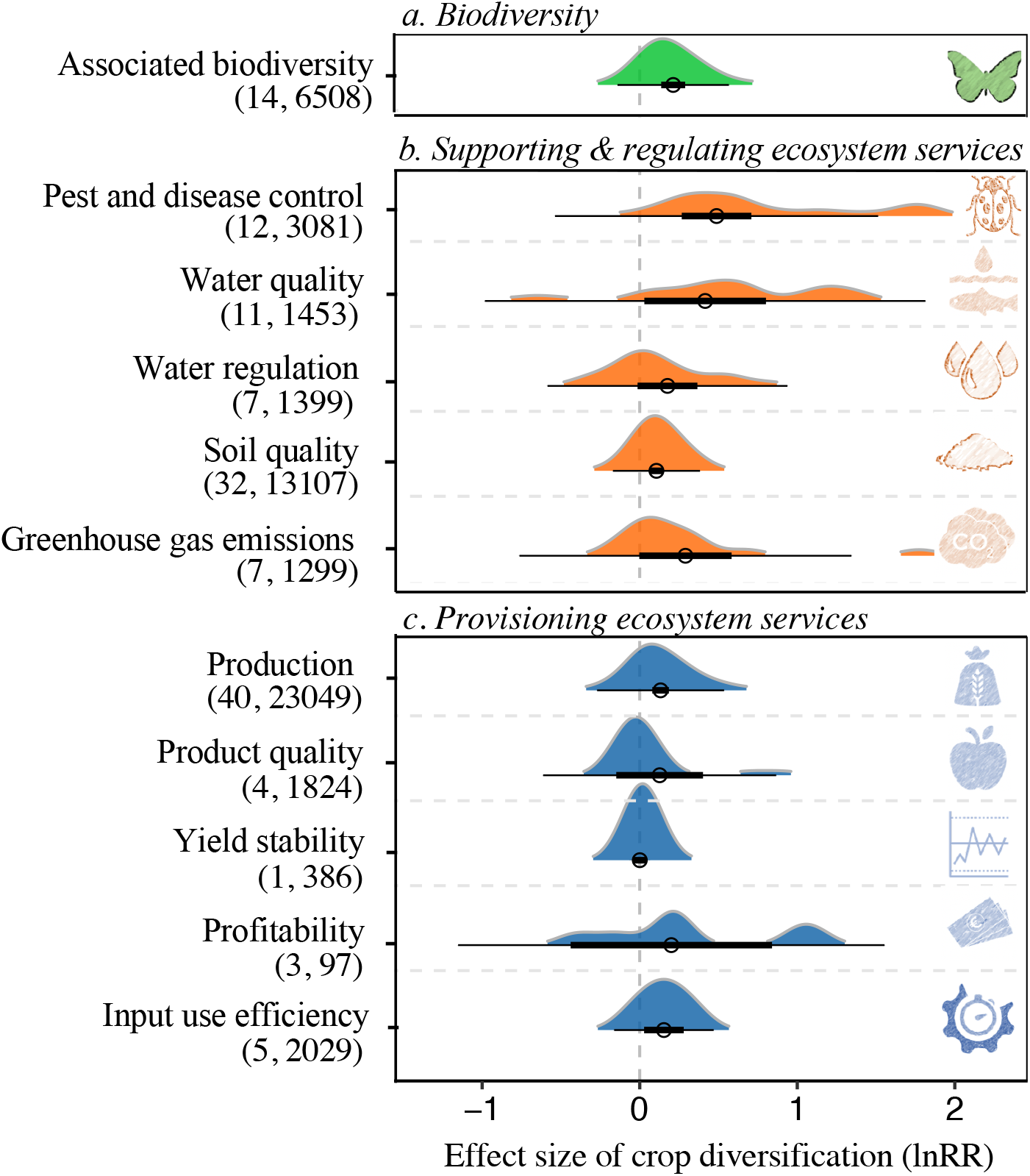
Impacts of crop diversification (all strategies) on associated biodiversity, supporting and regulating ecosystem services and provisioning ecosystem services. Points, thin lines and thick lines represent the estimated median summary effect, 95% confidence intervals (CI), and 95% prediction intervals (PI- see methods) of the logRatios estimated from a weighted mixed effect model. CIs indicate the range of values that likely contains the mean estimate of the data analysed, while PIs indicate the range of values that would likely contain the mean estimate if an additional new meta-analysis would be added. The number of meta-analyses and experiments are displayed between parentheses below each category. Colored shaded areas represent the distribution probability of the effect sizes of the meta-analyses.

The mechanisms behind the positive effect of crop diversification on associated biodiversity, regulating and supporting, as well as provisioning ecosystem services are manifold. Crop diversification can, for example, provide access to additional resources, or improve the diversity and connectivity of habitats for wildlife^13^. Regulatory services like pest and disease control, water or soil quality are, in turn, supported by the higher levels of associated biodiversity, as well as by changes in habitats and resource competition in more diversified systems. Globally, increasing crop diversity has, for example, a strong positive effect on all sub-categories of pest and disease control examined, i.e. weed reduction (60, CI:13%, 116%), pests reduction (33, CI: 2%, 74%), disease control (41, CI: 15%, 73%) and reduction of crop damages (62, CI: 37%, 93%). Increasing crop diversity could alter weed dynamics by limiting available resources (light, water, minerals) for weed species^26^. The increased canopy complexity of diversified systems also provides refuge to natural enemies^27,28^, reducing the spread of pests and diseases through microclimatic changes or physical barriers^29^. Similarly, crop diversification also improved all measured indicators of soil quality: soil physics, soil chemistry and soil carbon content (see supplementary Materials). The latter indicator is of particular interest to assess the potential for diversified systems to contribute to soil carbon sequestration and climate mitigation (although whether and to what degree agricultural practices can result in long-term carbon accumulation in soils is still debated^30,31^.

The higher levels of provisioning services observed in diversified systems could largely be driven by these positive effects on regulatory ecosystem services^32–35^. Garibaldi et al., 2018 showed that synergistic effects (positive interactions) between yield and regulatory services generally outnumber negative ones, suggesting that positive impacts of crop diversification on regulating services like soil quality and pest control lead to the positive impacts on provisioning services observed^36^.

The results of our analysis are, however, inconclusive for the regulating and supporting services of water regulation (+19%; CI: −1%, 44%) and GHG emissions (+25%; CI: 0%, 44%) despite thousands of experiments for these services (1400 and 1300 experiments, respectively). It is also important to note that GHG emissions is the only outcome examined that shows a negative trend – crop diversification practices thus might actually increase GHG emissions, but with results being non-significant due to high uncertainty and/or few syntheses. In addition, our literature synthesis, although exhaustive, shows that there is currently insufficient aggregated data to precisely evaluate the impacts of crop diversification on input use efficiency, yield stability, product quality and profitability – even though the tendency for these outcomes are also positive (i.e., improved input-use efficiency, and product quality, increased profitability). Our results are robust to the type of statistical model used and to publication bias (see Supplementary Materials).

### Different crop diversification strategies vary in the magnitude of their positive impacts

Despite these often-clear positive outcomes across all diversification practices, we observe high heterogeneity among the results of different meta-analyses (I2 statistics higher than 90% for most ecosystem services- Supplementary Materials), which implies that the effects and their magnitude depend on the crop diversification practice and/or bioclimatic conditions. Further analysis reveals that a substantial part of the variability of the mean effect sizes is due to significant differences in performances between the five crop diversification strategies.

The global positive effect of crop diversification on **associated biodiversity** strongly depends on the type of strategy considered (p-value < 0.001 – *Fig. 3; Supplementary Materials*). The biodiversity benefits are the largest for agroforestry (+61%, CI: 26, 105%), crop rotation (+37%; CI:16,62%) and cover crops (+21%, CI: 17, 25%)- which are all strategies that are often associated with heterogeneous non-crop habitats. Note that we also find substantial differences between different types of agroforestry practices (i.e., +9% of associated biodiversity for silvopasture; +45% for alley cropping; +62% for parklands, +86% for shaded perennial systems, see *Fig. 3*; for illustrative examples of these different agroforestry practices and for detailed definitions see Supplementary Materials). Intercropping and variety mixtures, on the other hand, have smaller or non-significant effects on associated biodiversity (+7%; CI: 3,12% and CI: 2; −12,18%, respectively).

**Figure 3.**
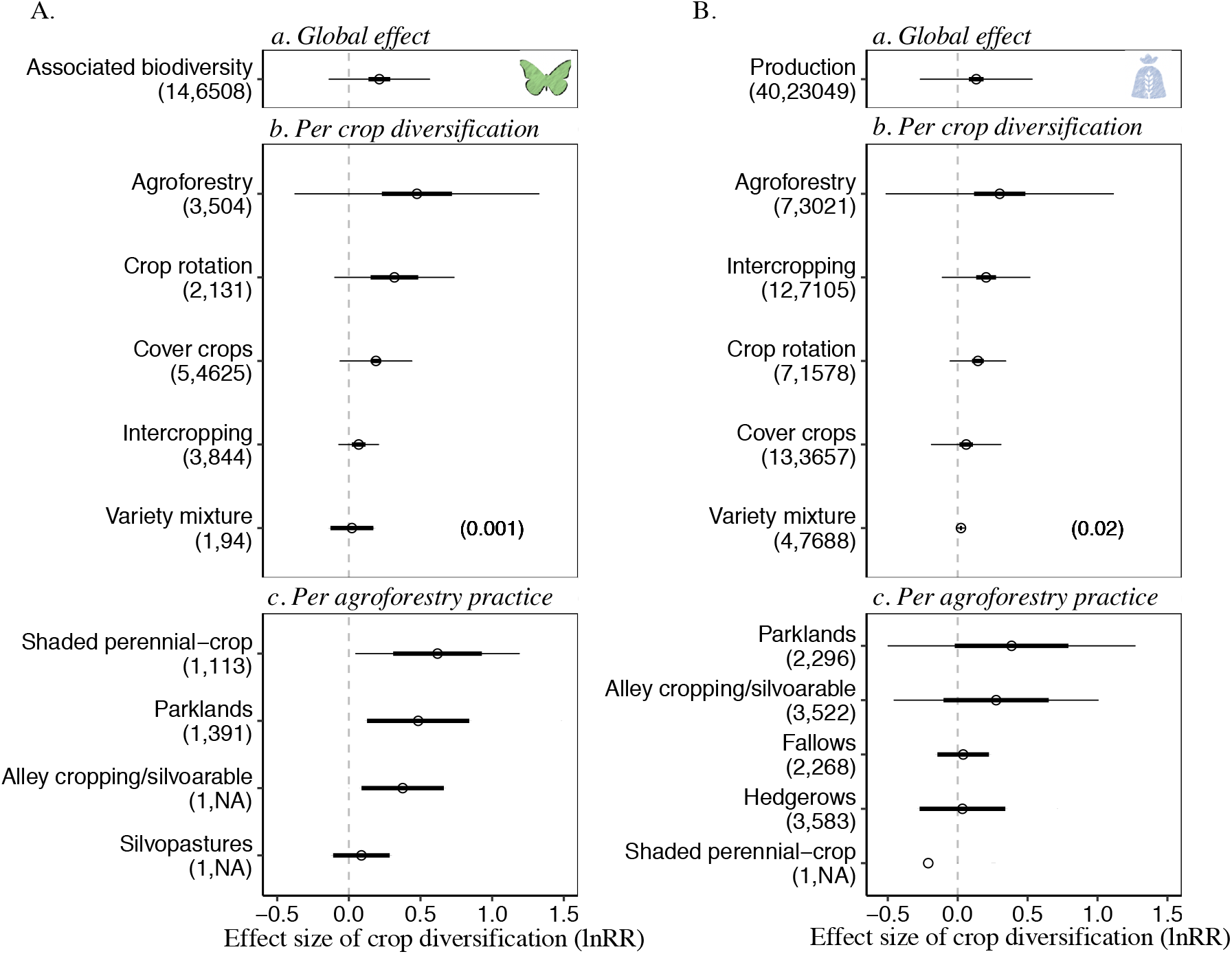
Impacts of the five crop diversification strategies on biodiversity (A.) and agricultural production (B.). Results are presented globally over all strategies (a.), for each of the five crop diversification strategy separately (b.), and for selected agroforestry practices (c.). The numbers of meta-analyses and experiments are displayed between parentheses below each category (NA indicates the absence of reported information the meta-analyses) on the left. Points, thin lines and thick lines represent the estimated median summary effect, 95% confidence intervals (CI), and 95% prediction intervals (PI) of the logRatios estimated from a weighted mixed effect model. The numbers between parentheses in sub-plots b. represent the p-values of statistical tests of differences between crop diversification strategies.

The variability of **agricultural production** observed for aggregated results is, also, largely explained by the different crop diversification strategies (67% of the variability of the effect sizes – p< 0.02 – *Fig. 3, Supplementary Materials*). The two diversification strategies relying on the simultaneous cultivation of different plant species within one field (i.e., agroforestry and intercropping) stand out as improving agricultural crop production most strongly (+35%; CI: 12%, 62%; +22%, CI: 14%, 32%, respectively; note that we do not consider tree production for agroforestry). Including an additional crop in a rotation leads to a 16% median increase (CI: 11%, 21%) in production levels of the following crops. Cover crops (+6%; CI: 1%, 12%) and variety mixtures (+2%; CI:1%, 3%) promote modest but still significant yield improvements compared to more simplified systems. The remaining variability of the effect sizes within each crop diversification strategy (reflected by the confidence intervals for each of the five strategies) could be caused by the diversity of conditions covered (e.g., a variety of agricultural practices, climates and soil characteristics, crop species or taxa, and initial diversification levels of the control plot). For agroforestry, for example, the different types of systems considered differed in their impact on agricultural production (i.e., −19% production for perennial shaded systems; +3% for hedgerows; +31% for alley cropping, +47% for parklands, see *Fig. 3*). For the other provisioning ecosystem services (i.e. product quality, input use efficiency, yield stability and profitability), the numbers of meta-analyses by crop diversification strategies are rather low, and do not allow a detailed comparison between strategies.

Most **supporting and regulating ecosystem services** also depend on the type of crop diversification strategy implemented. Pest and disease control is, for example, improved by all crop diversification strategies, but with different magnitudes of effects (*Fig. 4*). Our results suggest that the benefits are highest when using cover crops (+125%; CI: 83%, 178%), while intercropping (+66%; CI: 40%, 98%) and agroforestry (+59%; CI: 38%, 82%) still have high but slightly smaller positive impacts. For agroforestry, available evidence shows strong positive effects on pest and disease control for hedgerows (median effect = 84%) and shaded perennial systems (median effect = 40%; *Fig. 4*). The benefits of crop diversification for soil quality is also variable across strategies (p-value < 0.08- *Fig. 4*). The benefits are particularly important for agroforestry and intercropping (+19%; CI: 16%, 23% and +11%; CI: 5%, 18% respectively). Crop rotation promotes modest soil quality improvements (5%; CI: 2%, 8%), and variety mixtures have non-significant effects (−5, CI: −25%, 18%). For agroforestry, the median effect is −12% (non-significant) for silvopasture, +13% for hedgerows, +17% for alley cropping and +21% for parklands (*Fig. 4*). Other supporting and regulating ecosystem services such as water regulation (which includes variables like run-off reduction or soil water infiltration, Supplementary Material) also vary between strategies: Agroforestry leads to significant improvements (45%, CI: 13, 87%) while crop rotation and cover crops effects are non-significant (18%, CI: −5, 48% and 10, CI: −10, 34%, Supplementary Materials). On the contrary, no differences are observed among strategies for water quality, partly because of the very large estimated confidence intervals (89%, CI: 20, 200% for agroforestry; 87%, CI: 36, 156% for intercropping and 61, CI: 12, 132% for cover crops).

**Figure 4.**
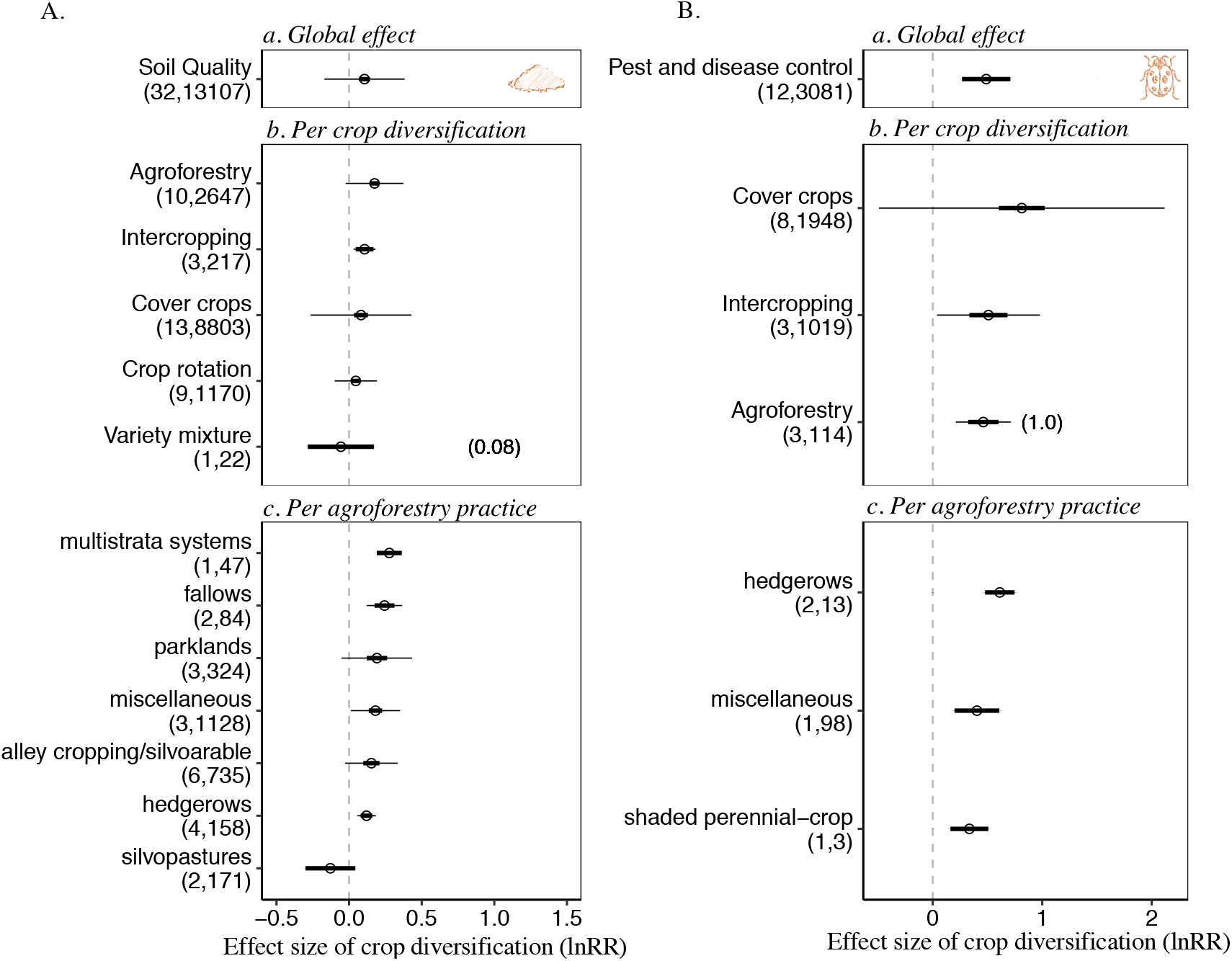
Impacts of the five crop diversification strategies on soil quality (A.) and pest and disease control (B.). Results are presented globally over all strategies (a.), for each of the five crop diversification strategy separately (b.), and selected agroforestry practices (c.). The numbers of meta-analyses and experiments are displayed between parentheses below each category (NA indicates the absence of reported information the meta-analyses) on the left. Points, thin lines and thick lines represent the estimated median summary effect, 95% confidence intervals (CI), and 95% prediction intervals (PI) of the logRatios estimated from a weighted mixed effect model. The numbers between parentheses in sub-plots b. represent the p-values of statistical tests of differences between crop diversification strategies.

### Climate impacts of crop diversification are variable and context-dependent

Given the importance of agriculture as a contributor to but also a potential solution to anthropogenic climate change^37^, any sustainable agricultural practice needs to be scrutinized for its climate mitigation potential. We find that agroforestry (+19%; CI: 14%, 24%), intercropping (+13%; CI: 6%, 10%) and cover crops (+13%; CI: 10%, 15%) contribute significantly to increased soil carbon content (see Supplementary Materials). On the other hand, both crop rotations and crop variety mixtures have a small and non-significant impact (+3%; CI: 0%, 4% and −5%; CI: −24%, 4% respectively). These different effects of different crop diversification strategies on soil carbon content may be linked with the fact that some strategies such as agroforestry, intercropping and cover crops tend to produce more above- and below-ground litter, which favors the formation and accumulation of soil organic carbon^38^ (see Supplementary Materials). This is in line with other recent studies that have highlighted the promises of agroforestry or cover crops for sequestering carbon in agricultural soils^39,40^. To contribute to soil carbon sequestration and climate mitigation, such practices need, not only provide additional biomass and carbon uptake but also (i) not result in a trade-off with yields (which would result in displacement effects), (ii) not result in the increase of other GHGs like nitrous oxide (N2O) or methane or increased energy use for example for fertilizer production, and (iii) persist over long periods^41,42^. While all diversification practices improve yields (see above), we find based on 6 meta-analyses that cover crops actually increase GHG emissions (+29%; CI: +49%, +1%). Experimental evidence suggests that the use of leguminous cover crops but also incorporation of cover crops into the soil, as well as the higher availability of mineralizable carbon under cover crop management can result in an increase of direct N2O emissions compared to non-cover crop management^43,44^. Yet, a literature synthesis assessing the net GHG balance of cover crops suggests that improved soil carbon stocks, as well as decreased indirect N2O emissions due to reduced nitrate leaching could outweigh the sometimes increased direct N2O emissions^44^. Another shortcoming of the use of cover crops for carbon sequestration is that cover crop management can be easily reversed and carbon stored released again in following years^41^. The net benefits of cover crops for climate mitigation are thus complex and appear to vary depending on factors like cover crop species, climate and tillage practices^41^.

More diversified agroforestry systems, instead, do not only improve yields (*Fig. 3*), result in more below-ground soil carbon (Supplementary materials), as well as more above-ground biomass (*Fig. 3*), but – being perennial systems – they are also likely to persist longer^45^ and some analyses suggest that they reduce other soil-based GHG emissions like N2O^46^ (which we were not able to assess due to a lack of agroforestry meta-analyses on GHG emissions). These important differences in the performances of different crop diversification practices for soil carbon and climate mitigation highlight the need to unpack such differences instead of treating crop diversification as a monolithic and always sustainable practice.

### Knowledge gaps and future research agenda

The transition to more sustainable diversified systems is a long and constantly evolving process requiring up-to-date and comprehensive multidisciplinary evaluation. The data gaps that we identified on some important outcomes like yield stability or farm profitability could partly explain the actual low rate of adoption of more diversified strategies in practice^47,48^. Most of the scientific literature on crop diversification has, to date, focused on ecological outcomes and production. Other agronomic outcomes are, instead, rarely studied. For example, few results are available on the economic performance of crop diversification strategies (*Fig. 2*), certainly a major factor limiting the rate of adoption of any new agricultural practice^47^. Restoring ecosystem services or substituting ecosystem functions provided by petrochemical energy will only likely be realized by farmers and land users if there is a direct economic or other benefit to be derived from it^8^. Similarly, evidence on the stability of the production of diversified crop management is scarce (only one meta-analysis, *Fig. 2*), despite the abundant theoretical literature that suggests d that diversified agro-ecosystems might be better able to cope with perturbations (e.g. ^5,48^). More scientific evidence on the impacts of crop diversity on product quality (e.g. micronutrient content) and input use efficiency – also two variables of interests to farmers and consumers – could also favor a wider adoption.

Beyond the field scale that constitutes the basis of this study, it is also important to maintain ecosystem services at larger scales, such as farm or landscape, and to promote a higher diversity of land-uses (e.g.^49^). Most of the ecological processes are strongly dependent on the complex interactions between land use, biophysical context and human activity at the landscape scale, such as water quantity and quality, pollination, pest regulation or carbon storage^50,51^.

## Conclusion

Our results, based on the largest assessment examining effects of increasing the diversity of cultivated crop species or varieties in agroecosystems, show that these effects are positive but variable and depend on the type of crop diversification strategy considered. Our analysis is based on hundreds or even thousands of field-scale experiments where the effects of increased crop diversity have been compared to less diverse agrosystems. Our synthesis allows the identification of strategies that effectively and consistently reduce air, soil and water pollution and increase biodiversity, pest and diseases control and crop production. Agroforestry strikes out as a particularly promising strategy that is able to substantially increase all the ecosystem services considered in our analysis (and for which sufficient data was available to allow drawing robust conclusions), i.e. associated biodiversity, production, water regulation, water quality, pest and diseases control and soil quality. The other diversification strategies also have important effects on specific ecosystem services, e.g. cover crops and intercropping on water quality and pests and disease control or crop rotations on associated biodiversity and production. Variety mixtures, instead, provide the lowest benefits for most of the ecosystem services considered.

Despite a sometimes-uneven geographic repartition of available studies^19^, our exhaustive and quantitative review of the literature, by precisely weighing the evidence based on its quality, redundancy, and precisely analyzing its heterogeneity, provides a solid basis for policy makers and scientists alike to build future decision-making and future research upon the existing knowledge base. It can also contribute to help the Food agricultural Organization (FAO) monitoring and orienting policies towards Sustainable Development Goals or provide support for the new European “Green Deal” as a starting point for regional debates on the effectiveness of the various strategies analyzed globally^52^. These broad conclusions should, however, not hide the fact that heterogeneity exists and that the results of a single meta-analysis could sometimes diverge from our mean estimates, as indicated by the wide prediction intervals found in our analysis. Agro-ecological transitions must therefore be designed to take into account local contexts and constraints^53^

## Methods

### Data collection and extraction

We identified gray and published meta-analyses on the impacts of crop diversity on biodiversity and ecosystem services through a systematic search using Web of Science, CAB abstract, Greenfile, Environment Complete Database, Agricola and Google Scholar. The search was performed in May 2020 and updated in November 2020 with the equation: (*meta-analysis OR meta analysis) AND (cropping system OR crop* OR agriculture) AND ((rotation OR Diversification OR intercrop* OR cover crop OR mixture) OR (organic AND (system OR agriculture)) OR (conservation AND (system OR agriculture)) OR no till* OR agroforestry OR agroecology*). The query was carried out on the article title, abstract and keywords, with no restriction on the date of publication. We also screened the references cited in each selected meta-analysis. This resulted in a total of 984 publications (844 once duplicates removed – see Supplementary Materials). Only publications satisfying the following criteria were retained for the analysis (i) the publication reports the results of a quantitative analysis based on several primary studies, i.e. a meta-analysis, (ii) The meta-analysis focus on at least one crop diversification strategies defined in table S1, (iii) The adjacent control plot used in the meta-analysis is less diversified as the treatment (we exclude comparisons to diversified natural ecosystems), (iv) the meta-analysis present indicators of variability of the effect-sizes. Studies dealing with pure forestry or wood production were excluded. Based on these criteria, 95 meta-analyses were selected.

All quantitative measures of the effects of crop diversity compared to the less diversified reference, i.e effect-sizes, were extracted from text, table or figure using Plot Digitizer (http://plotdigitizer.sourceforge.net/). The effect-sizes of each meta-analysis were precisely documented (unit, name of the variable, soil, climate, management conditions, etc.). We also retrieved the number of primary studies that were used to calculate the effect-sizes, the number of experiments (one primary study can provide several independent estimates), and the confidence intervals or other indicators of dispersion, when available. Effect sizes expressed as standardized mean difference or percent changes were then converted into ratios, to ensure the comparability of the results between the meta-analyses^54^. Different meta-analyses could be partly based on the same pool of individual experiments. To handle this pseudo replicated data for our second-order meta-analysis, we retrieved all the references of primary studies of each meta-analysis (see Supplementary Materials). We then calculated a redundancy index between each pair of meta-analyses, and included it in our meta-analytical model (see *DataAnalysis section*).

The meta-analyses differ in the quality of the methodology used to retrieve the primary study and analyze the data^19,55^ which can result in biased and misleading results^56^. To lower the importance of low-quality studies in our second-order meta-analysis, we calculated a quality score, based on 20 criteria analyzing the literature review, statistical analyses and analysis of potential bias of each meta-analysis (see Supplementary Materials). We then integrate this score as a proxy of the internal quality of the meta-analyses in our meta-analytical model (see *Data Analysis section*).

All retrieved data are freely available on internet web platform: https://cropdiversification.shinyapps.io/Crop_diversification_2020

### Data analysis

We categorized the retrieved response variables reported in the meta-analyses into main- and sub-categories of biodiversity, supporting and regulating ecosystem services and provisioning ecosystem services. The 12 main categories are associated biodiversity, pest and disease control, water quality, water regulation, soil quality, greenhouse gas (GHG) emissions, agricultural production, product quality, yield stability, profitability and input use efficiency. Sub-categories are shown in Supplementary Materials.

We quantified the impacts of crop diversity on each main and sub-categories of biodiversity, ecosystem services. We then detailed the impacts for each crop diversification strategy (e.g. agroforestry, intercropping). The global form of our models is:

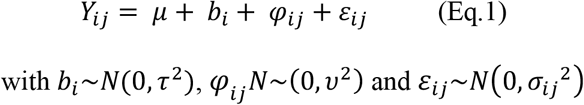

Where *Y_ij_* are the j^th^ effect-size of the i^th^ a meta-analysis, *μ* is the mean effect, *b_i_* is the random meta-analysis effect, *φ_ij_* is the random effect-size effect within the ith meta-analysis, and *í_ij_* is the random estimation errors associated with the j^th^ effect size of the i^th^ meta-analysis. We refer to *τ^2^* as the between-meta-analysis variance, *ν^2^* as the between effect-size variance, and *σ_ij_^2^* as the variance of the j^th^ estimated effect-size of the i^th^ meta-analysis.

We weighted each meta-analysis by the inverse of the variance of the effect-size, as recommended by ^57^. We then reduced the weight of the studies of lower quality following doi et al., 2008 methodology^58^. We considered the non-independence between the effect sizes of different meta-analyses by calculating variance-covariance matrix based on a pseudo correlation between meta-analyses^59^. The proxy of the correlation between each pair of meta-analyses was estimated as, where m is the number of common primary studies between each pair of meta-analyses, and n and n2 are the total number of primary studies in the two meta-analyses, respectively.

We analyzed the heterogeneity of the estimated effect-sizes by calculating i) the 95% confidence-intervals, i.e. the range of values that likely contains the mean estimates, ii) the 95% prediction intervals, i.e. the range that likely contains the mean value of a single new observation, a new meta-analysis in our case, iii) the I2 statistics, i.e. the percentage of variation across studies that is due to heterogeneity rather than chance^60^. We also precised the percentage of variability of each biodiversity and ecosystem services explained by the crop diversification strategy. The added-value of adding crop diversification strategy the models was assessed by a likelihood ratio test between model including crop diversification as moderators and a null model.

The potential publication biases were assessed with funnel plots and egger tests^61^. The funnel plots assume that studies with high precision will be plotted near the average mean effect, and studies with low precision will be spread evenly on both sides. The Egger value tests for the asymmetry of the funnel plot.

We also test the robustness of our results to the type of model used (with or without random effect), the type of random effects included in the model (ID meta-analysis and scenario vs. only ID meta-analyses), and the potential impact of quality of the results (model considering quality of the meta-analyses or not). The sensitivity of the results against publication bias was tested with Rosenthal fail-safe number, i.e. the number of additional studies with a mean null result necessary to reduce the nullify the global estimated effects (see Supplementary Materials). We also provide an estimation of the mean effect considering the missing studies, based on trim-and-fill methodology^62^.

The parameters of the model were estimated by restricted maximum likelihood. We used log-transformed ratios for performing the statistical analyses. All statistical analyses were conducted with R^45^ (version 3.0.2), dplyr^63^ for data management, metafor for model analysis^64^ and ggplot2 for data visualization^65^.

## Contribution

DM, TBA and DB participated in the design and the coordination of the study. DB performs the data collection, with the help of DM and TBA. DB performed the statistical analyses with the help of DM, and designed the figures. DB and TBA wrote the draft, all authors provided critical feedback and contributed significantly to the writing.

## Acknowledgments

This work was produced within the framework of the European project ‘Diversification through Rotation, Intercropping, Multiple Cropping, Promoted with Actors and value-Chains towards Sustainability’ (DiverIMPACTS), funded by the European Commission under Grant Agreement number 727482. It was also supported by the INRAE-CIRAD metaprogram GloFoods and by the Institute of Convergence CLAND (16-CONV-0003). We are grateful to Mathilde Duvallet for her contribution to the database.

## Data availability statement

The data that support the findings of this study are openly available on an internet web platform: https://cropdiversification.shinyapps.io/Crop_diversification_2020

## Supplementary materials

### Identification and selection of the studies

**Figure S1.**
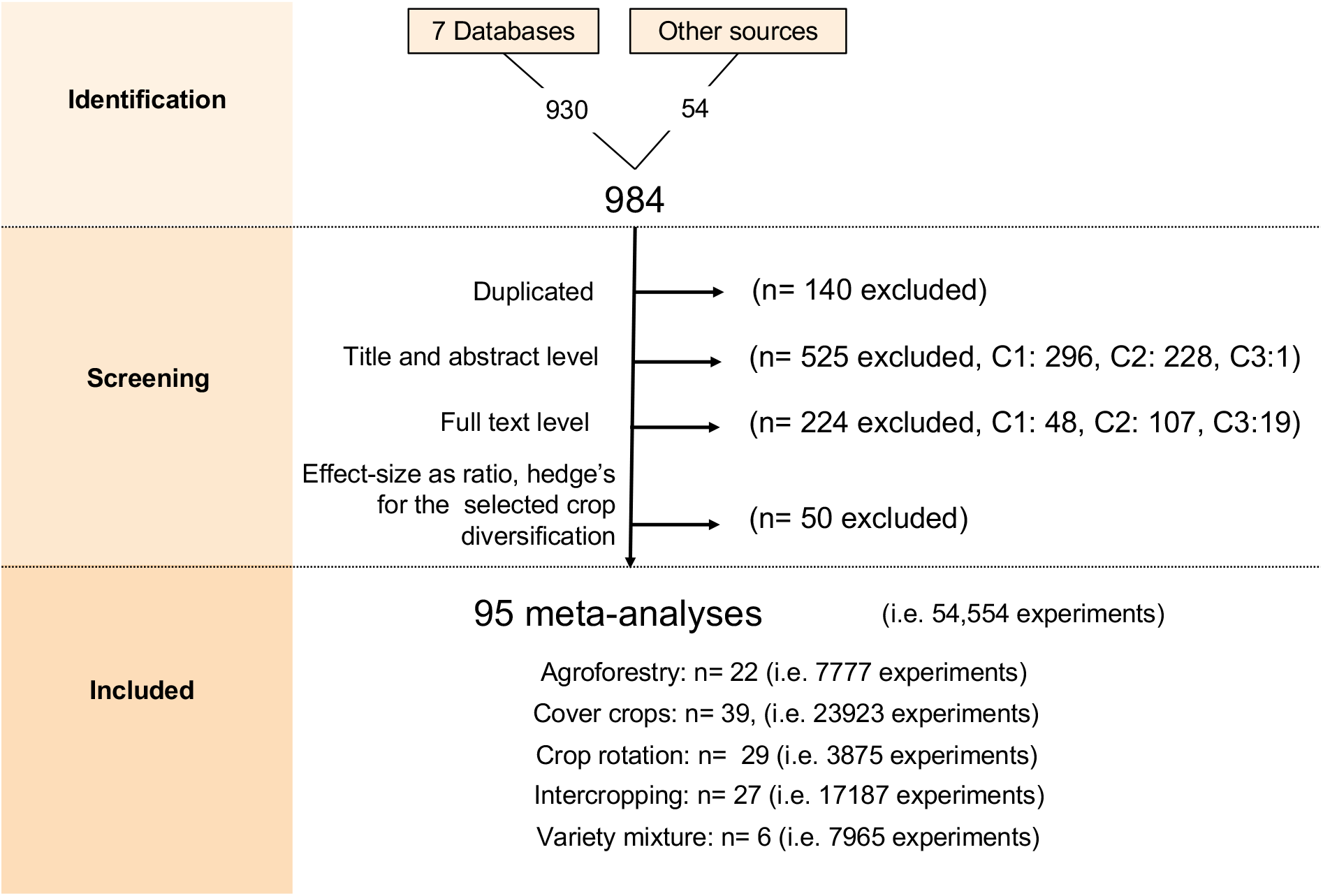
Prisma flowchart of literature search and screening process. C1: Selection criterion 1 (several individual studies are analyzed); C2: Selection criterion 2 (assessment of the impact of at least one strategy of crop diversification). C3: Selection criterion 3 (control plots are present next to treatment plots)

### Definition of the crop diversification considered and illustrative examples

**Table S1.**
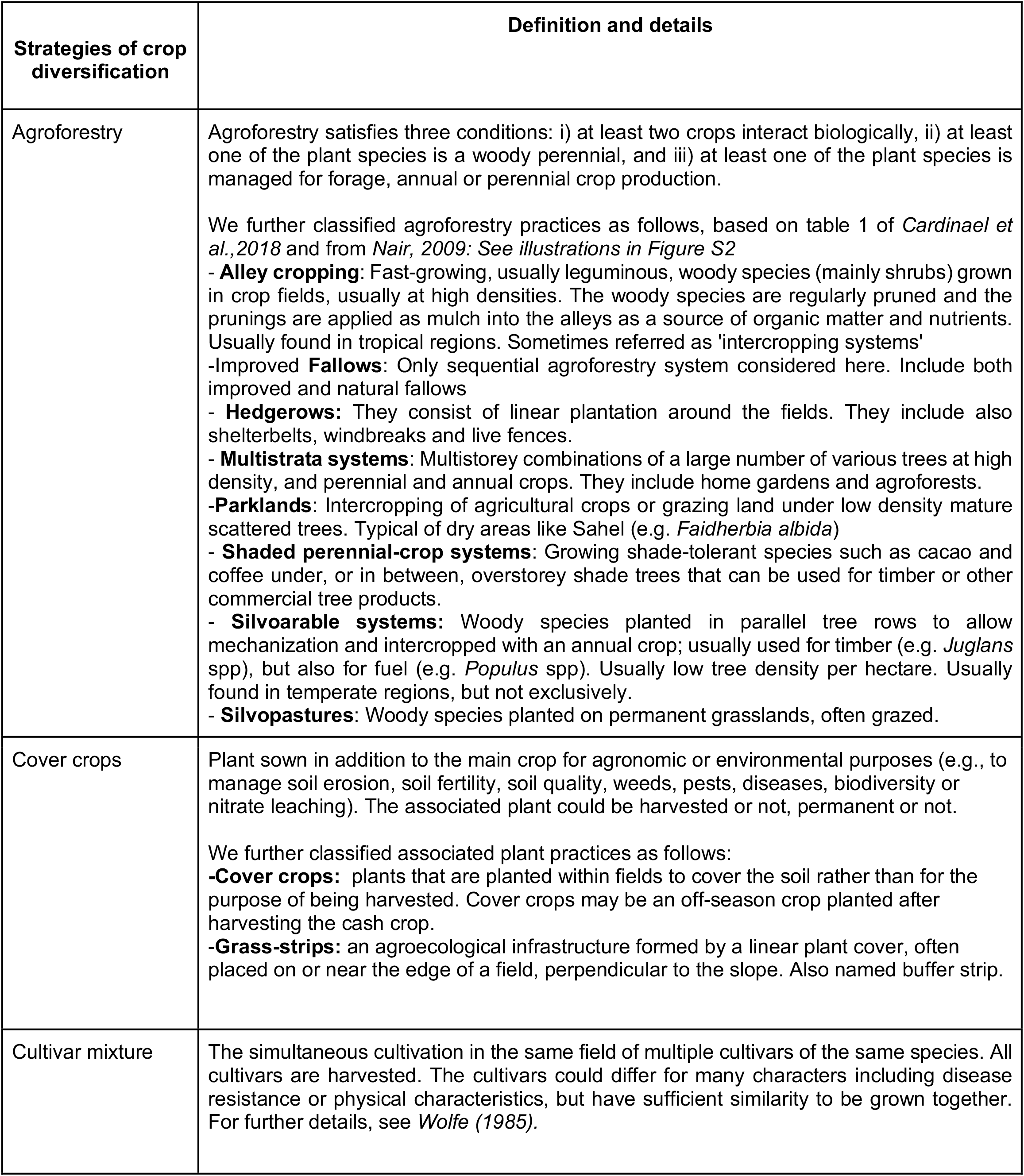
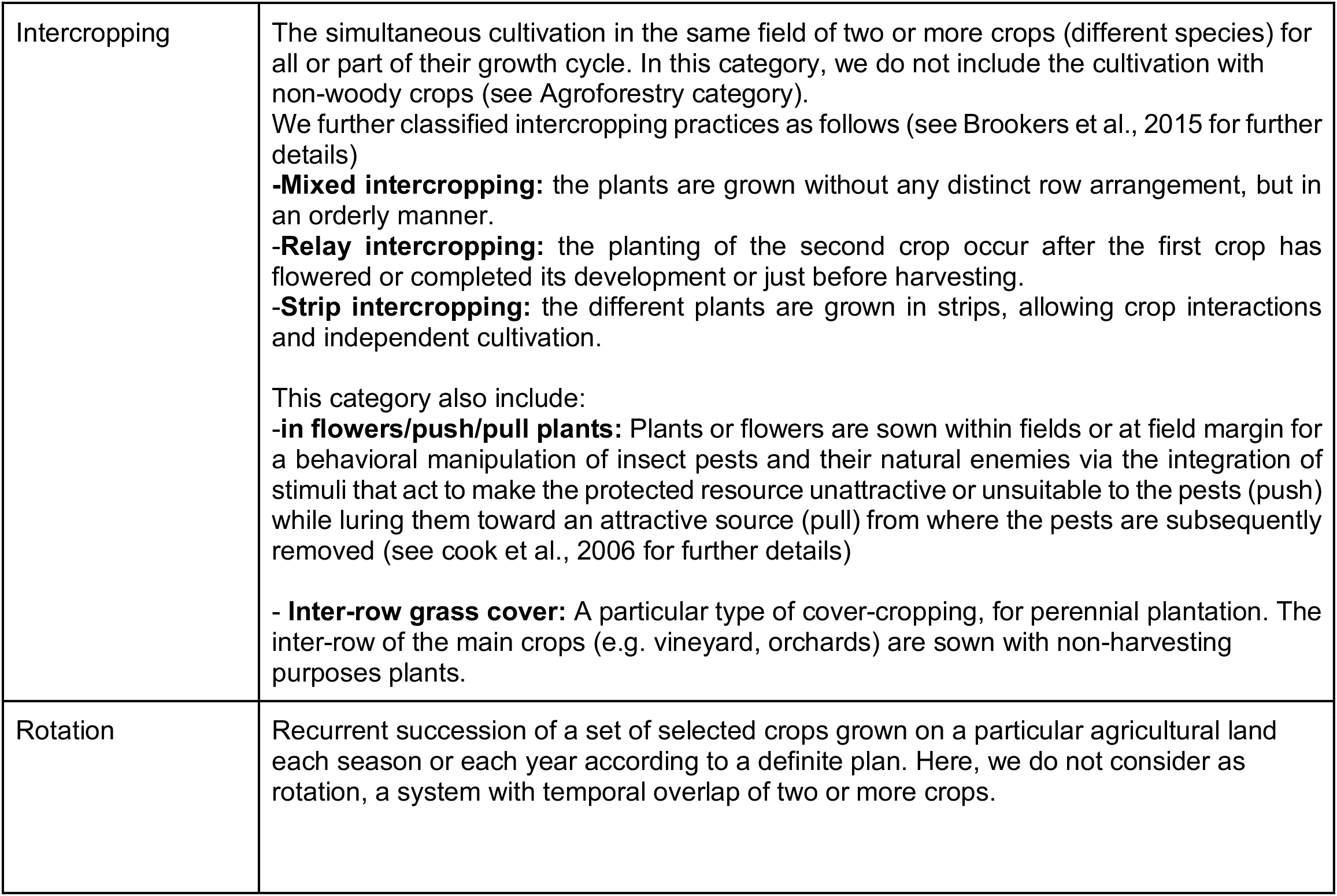
Definition of the crop diversification. considered

**Figure S2.**
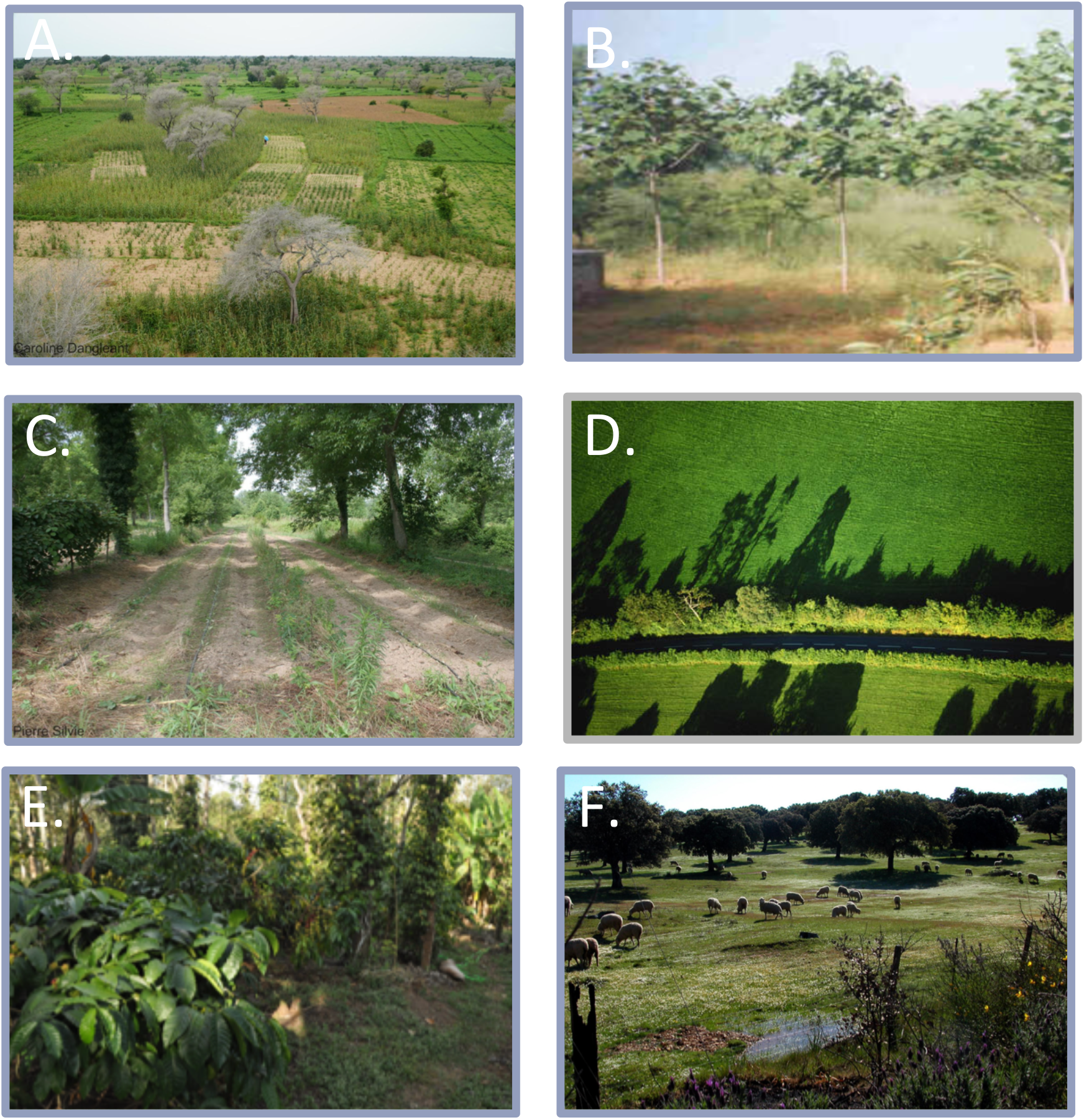
Illustrative example of the six types of agroforestry practices considered in our study. A. Parkland (Faidherbia albida, Pennisetum glaucum and Acacia albida, Senegal); B. Improved fallows (Gmelina_arborea); C. Alley cropping/silvoarable (Lactuca sativa under *Juglans regia* L., France), D. Hedgerows (England), E. shaded perennial crops (Coffea arabica, India), F. Silvopasture (dehesa, Spain).

Text1. Photos credits.

### Further details on the effects-sizes, and their variability

**Figure S3._**Percent of intra-(blue) and the sum of intra-and inter- (orange) variability explained by diversification strategies for different biodiversity and ecosystem services. The numbers at the top of the plot represent the p-values of the test of difference between diversification strategies.

**Table S2.** Global description of available data presented as ratio and their characteristics. Rosenthal’s Fail-Safe Number represent the number of additional ‘negative’ studies (studies in which the intervention effect was zero) that would be needed to increase the p value for the meta-analysis to above 0.05 (*Rosenthal 1979*). The heterogeneity (I2) represent the percentage of the variability in effect estimates that is due to heterogeneity rather than sampling error. The proportion of total variance that can be attributable to within-cluster heterogeneity is indicated in the last column.

| **Ecosystem services** | **number of**<br>**primary meta-**<br>**analyses** | **number of**<br>**effect_sizes** | **number of**<br>**paired -data** | **Rosenthal's**<br>**Fail-Safe**<br>**Number** | **I2** |
| --- | --- | --- | --- | --- | --- |
| Yield stability | 1 | 4 | 366 | NA | NA |
| Profitability | 3 | 7 | 97 | 456 | 88,20 |
| Product quality | 4 | 21 | 1824 | 36 | 99,84 |
| Input use efficiency | 5 | 10 | 2029 | 2393 | 98,58 |
| Greenhouse gas emissions | 7 | 13 | 1299 | 7257 | 99,39 |
| Water regulation | 7 | 22 | 1399 | 3423 | 98,76 |
| Water quality | 11 | 33 | 1453 | 6722 | 98,5 |
| Associated biodiversity | 14 | 100 | 6508 | 63928 | 99,37 |
| Pest and disease control | 12 | 59 | 3081 | 17866 | 94,87 |
| Soil quality | 32 | 151 | 13107 | 347734 | 99,92 |
| Production | 40 | 122 | 23049 | 101676 | 99,92 |

**Table S3.**
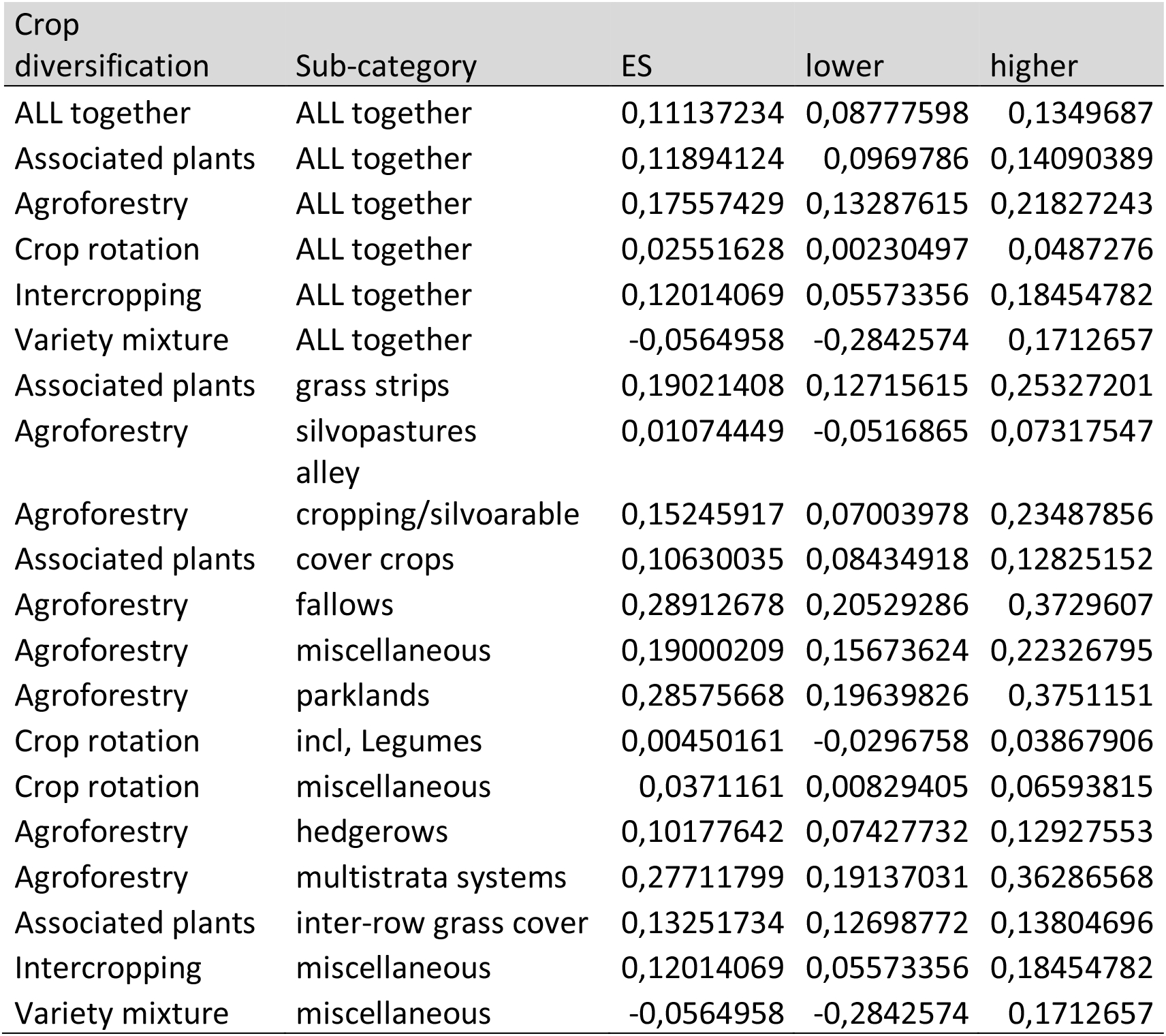
**Detailed effect-sizes for s**oil carbon (sub-category of soil quality). The effects-sizes are presented in logRatio for main and sub-categories of crop diversification. **All effect-sizes are available in the web app: https://cropdiversification.shinyapps.io/Crop_diversification_2020**

**Figure S4.**
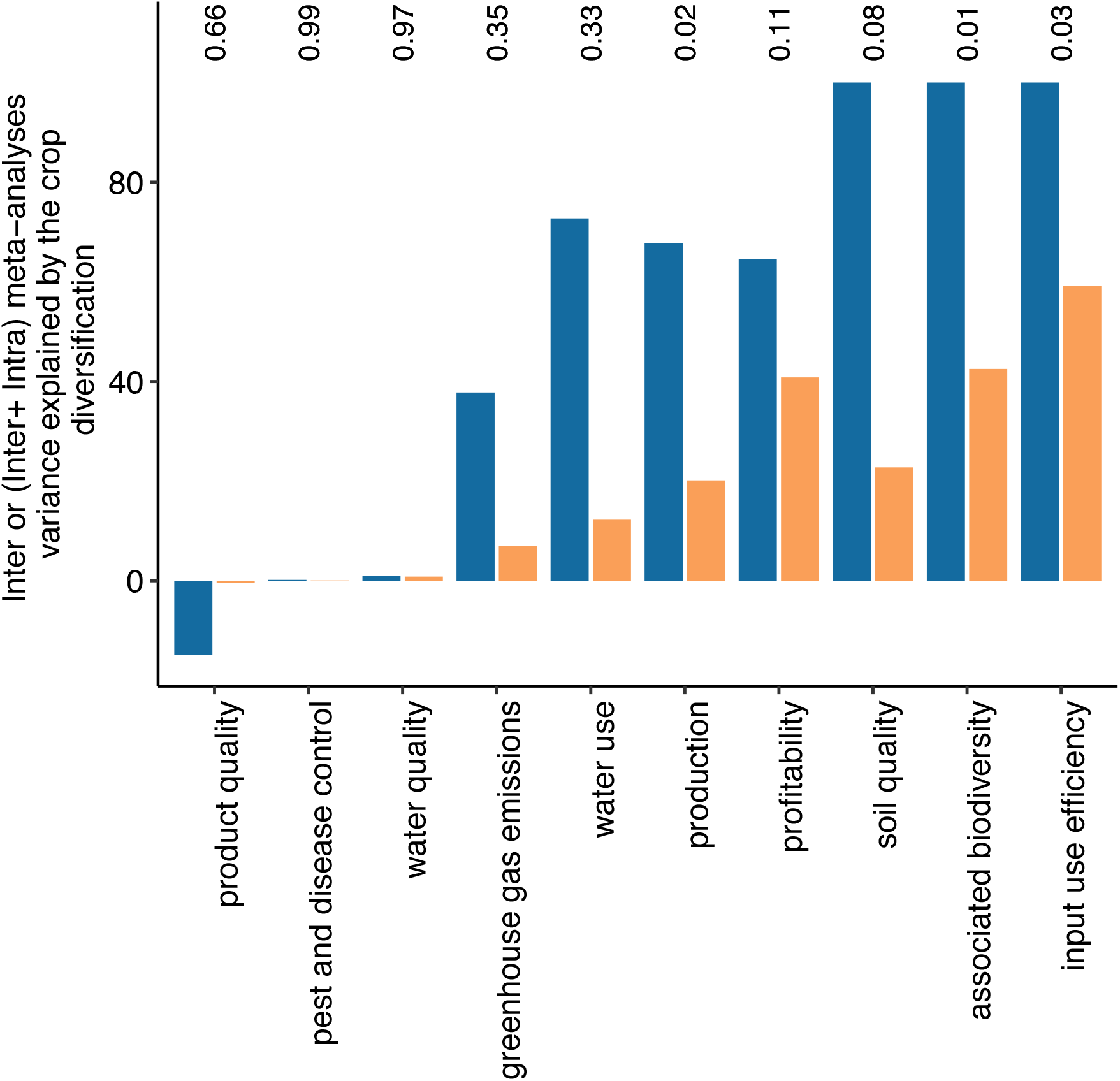
Description of individual variables considered within each Main categories of biodiversity and ecosystem services,

### Quality of the included meta-analyses

**Table S5.** **Description of the 20 individual quality items**

| Item | description |
| --- | --- |
| A protocol is published before the meta-analysis | An a priori review protocol exists, and can be accessed (e.g., Web address), and, if available, provide registration information including registration number. |
| The aim of the study is clearly mentioned | The list of criteria used to defined the scope of the study are clearly mentioned PICO : Population, variable response, treatment and control. |
| Literature databases are mentioned | The study describes the databases in the search. |
| Additional literature search is performed | Additional research (grey literature?) |
| Search strings are presented | Are the search string clearly defined? All search terms, so that the exact search is repeatable by a third party |
| Inclusion and exclusion criteria are mentioned | Description of the criteria used to select articles from the original result list (to ensure repeatability). If additional exclusion motives - these are also detailed (e.g., years considered, language, publication status, date last searched) |
| The number of studies is given at each selection steps | Detailed summary of studies screened, assessed for eligibility, and included in the review, ideally with a flow diagram. Diagram PRISMA |
| The list of included studies is provided | The study includes the final list of the references of the selected individual studies used in the meta-analysis |
| The list of excluded studies is provided | The study includes the list of the excluded studies |
| Method of data extraction is described | Do the authors explicate how data were extracted from the figures? |
| The dataset is available | Availability of the original dataset? (In results?) |
| All tools used for the review are mentioned | Name of the program/package used for statistical analysis |
| Statistical models are described | Statistical analysis of the database: description (model, effect(s), equation...) |
| Studies are weighted according to their accuracy | Are individual studies weighted according to their accuracy or quality? |
| Effect sizes of individual studies are presented | Provide information of the size effect for each individual study includes in the meta-analysis? (table, plot) |
| Confidence intervals of individual studies are mentioned | Provide information of the confidence interval for each individual study included in the meta-analysis? (table, plot) |
| Quantitative results of the meta-analyses are fully described | Results: Quantitative information on summary effects (effect estimates and confidence intervals, ideally with a forest plot). |
| Heterogeneity of results is analysed | Is the heterogeneity analyzed using sensitivity analysis or meta-regression? |
| Publication bias is analysed | Does the synthesis consider possible publication bias? |
| Funding sources are mentioned | Describe sources of funding for the systematic review and other support (e.g., supply of data); role of funders for the systematic review. |

**Figure S5.**
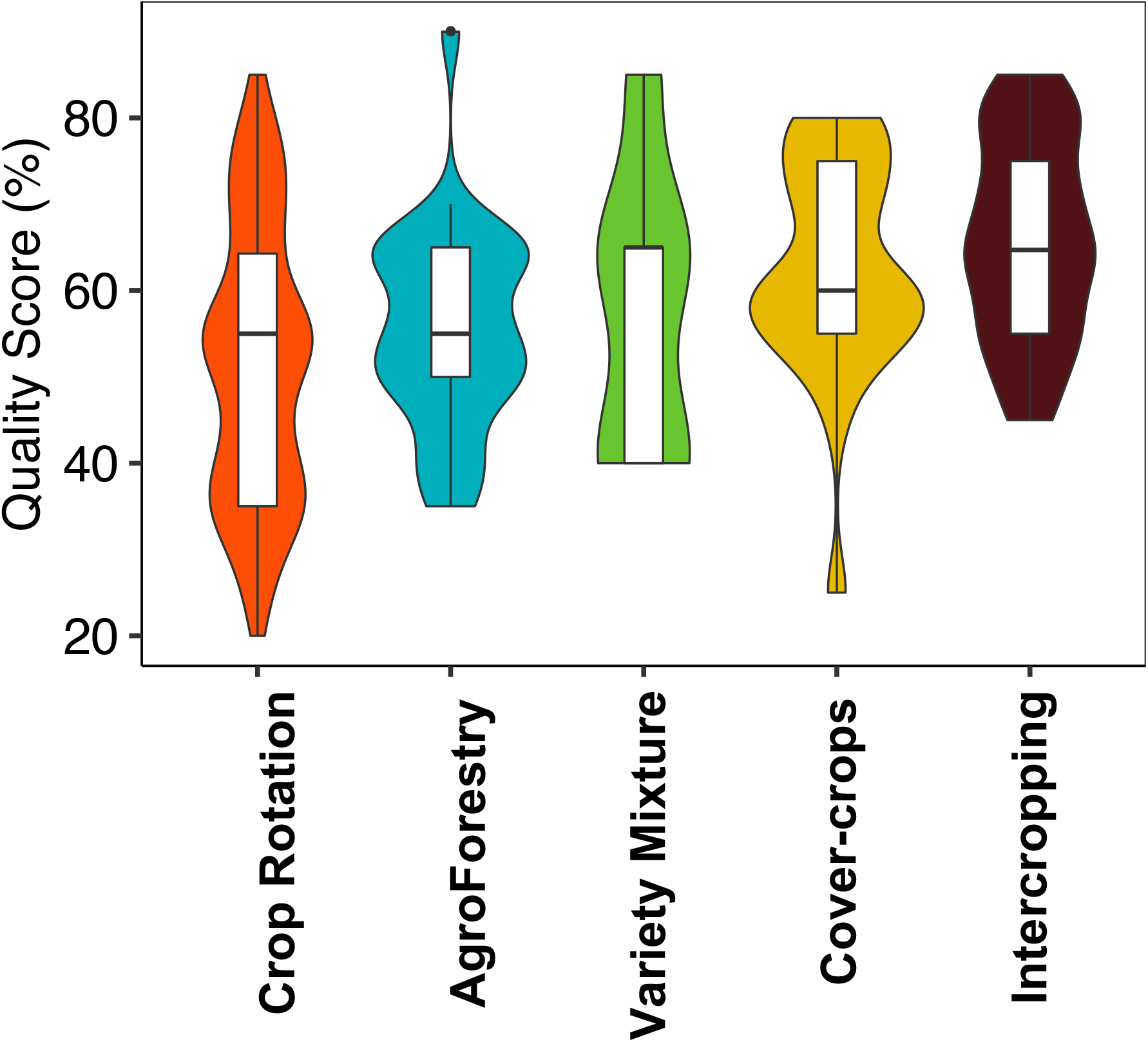
Boxplot of the quality score for each crop diversification strategies. The quality score represents the proportion of individual quality criterions met. Precise description of quality items are available in Table S3.

**Figure S6.**
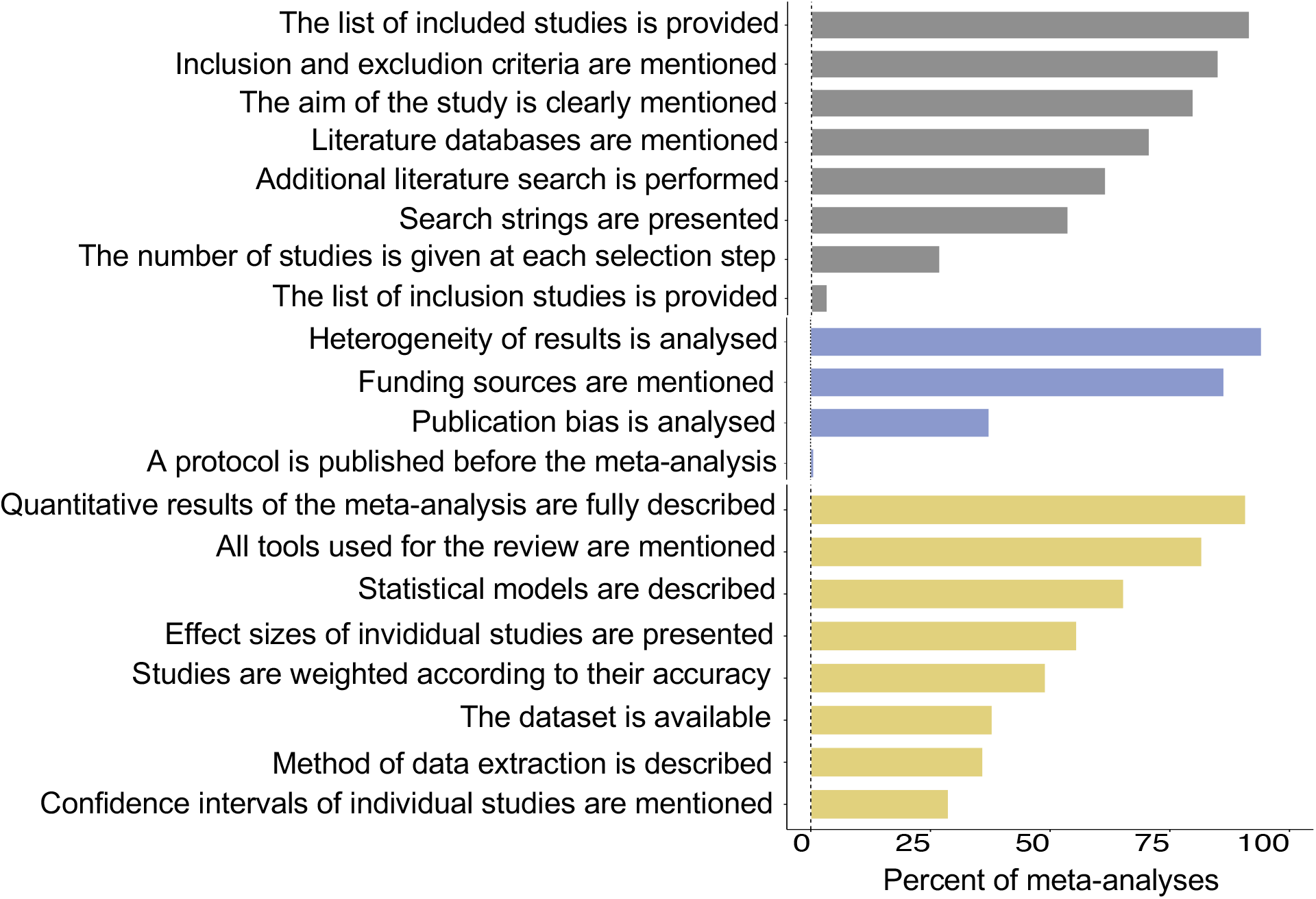
Percentage of meta-analyses meeting the 20 individual quality criteria. Quality criteria are organized in three main groups: review criteria (grey bars), statistical analyses (yellow bars) and bias (blue bars). Precise description of quality items are available in Table S3.

### Redundancy of the included meta-analyses

**Figure S7. Apparent redundancy of primary studies among meta-analyses**. Results on the x-axis are expressed as fractions of common primary studies in two meta-analyses.

### Analysis of the robustness of our conclusion

**Figure S8.**
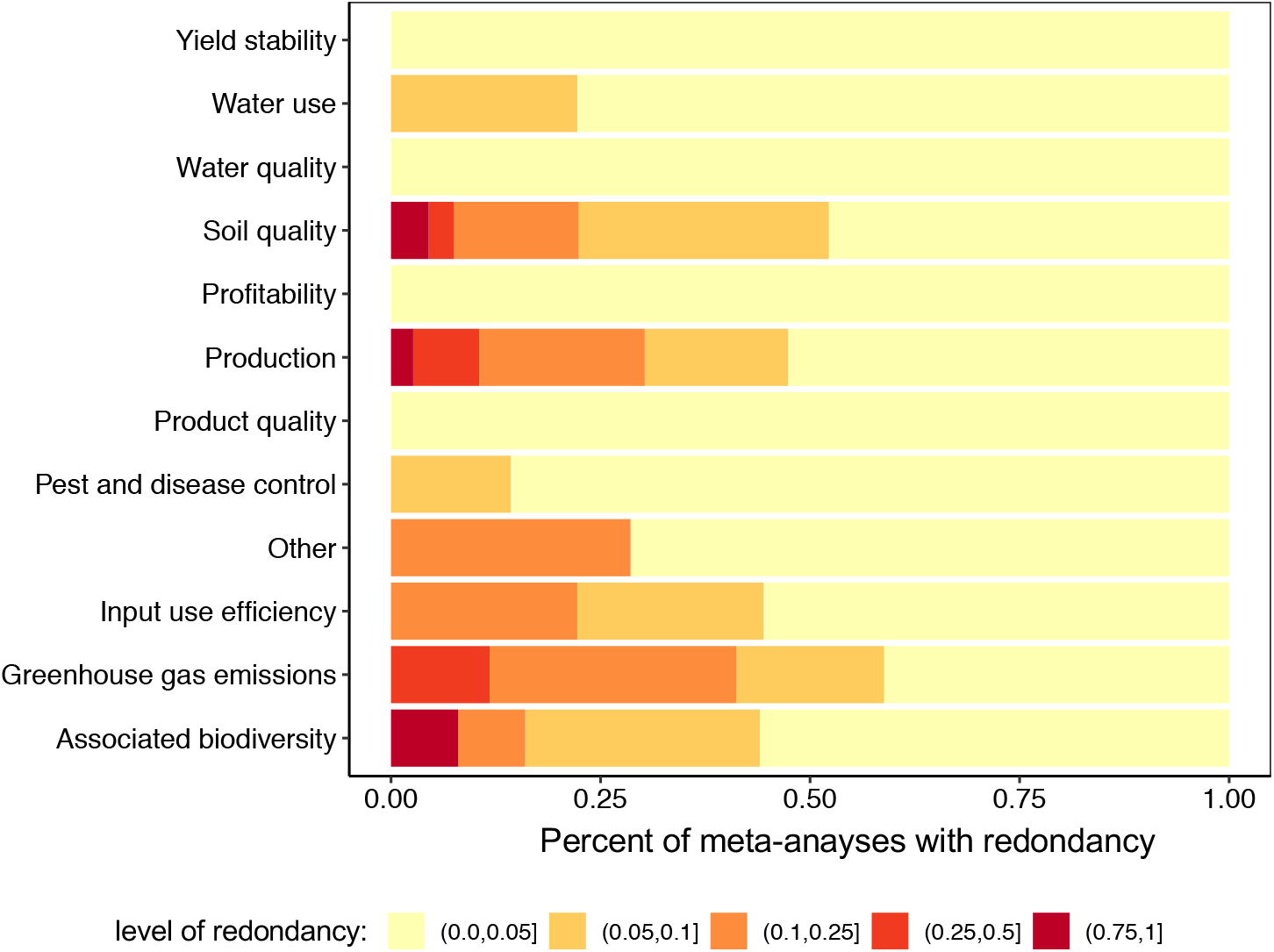
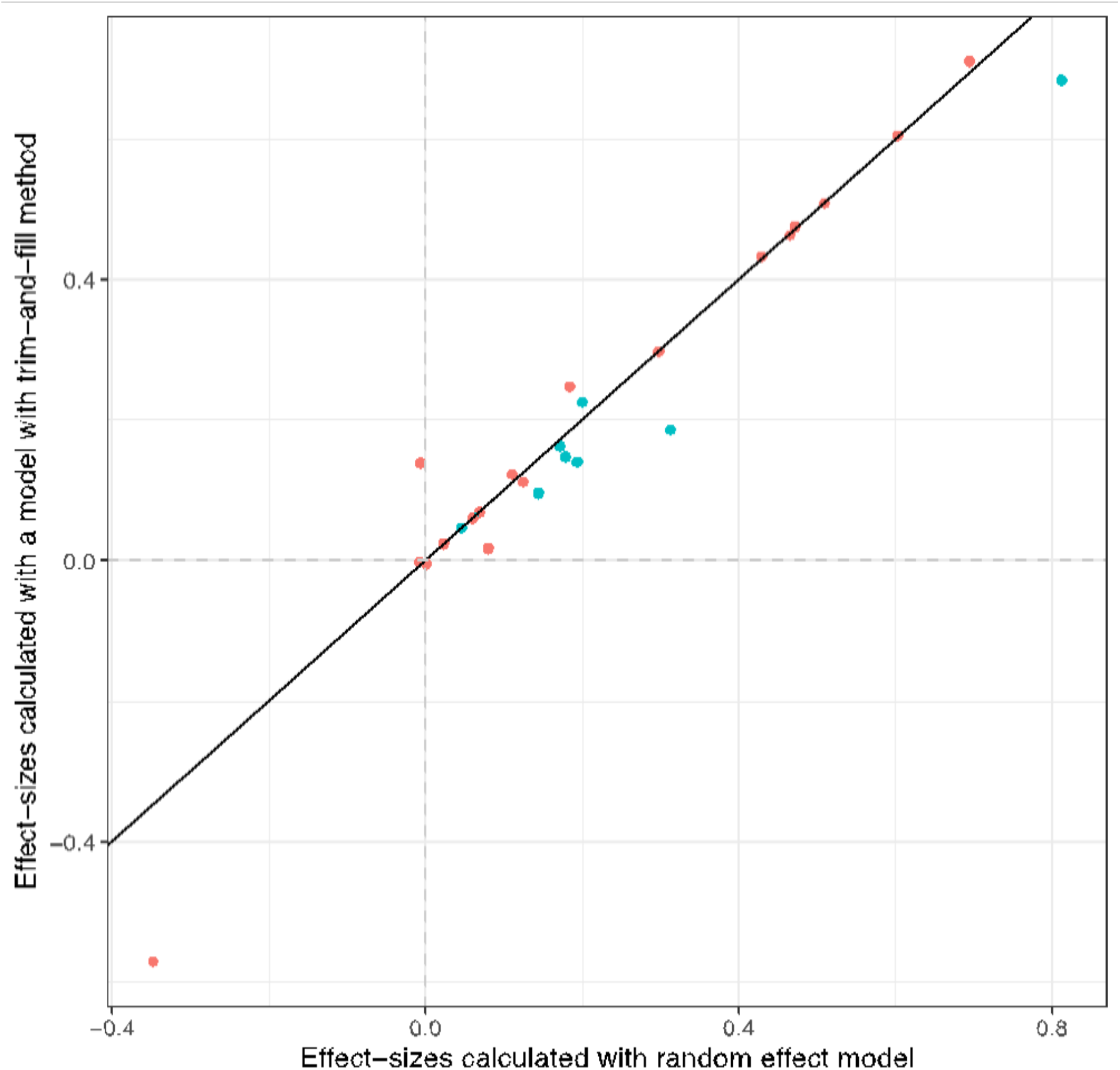
Analysis of the impact of potential publication bias. The mean effect-sizes calculated with a random effect model (ID and scenario as random effects) are plotted against the same effect-sizes calculated with the trim and fill method (see *Duval et al., 2000* for further details). Blue points indicate significant egger tests (p-value < 0.005). Results are presented for the five strategies of crop diversification on the main ecosystem services. For details on the few categories presenting significant egger test, see Table S2

**Figure S9.**
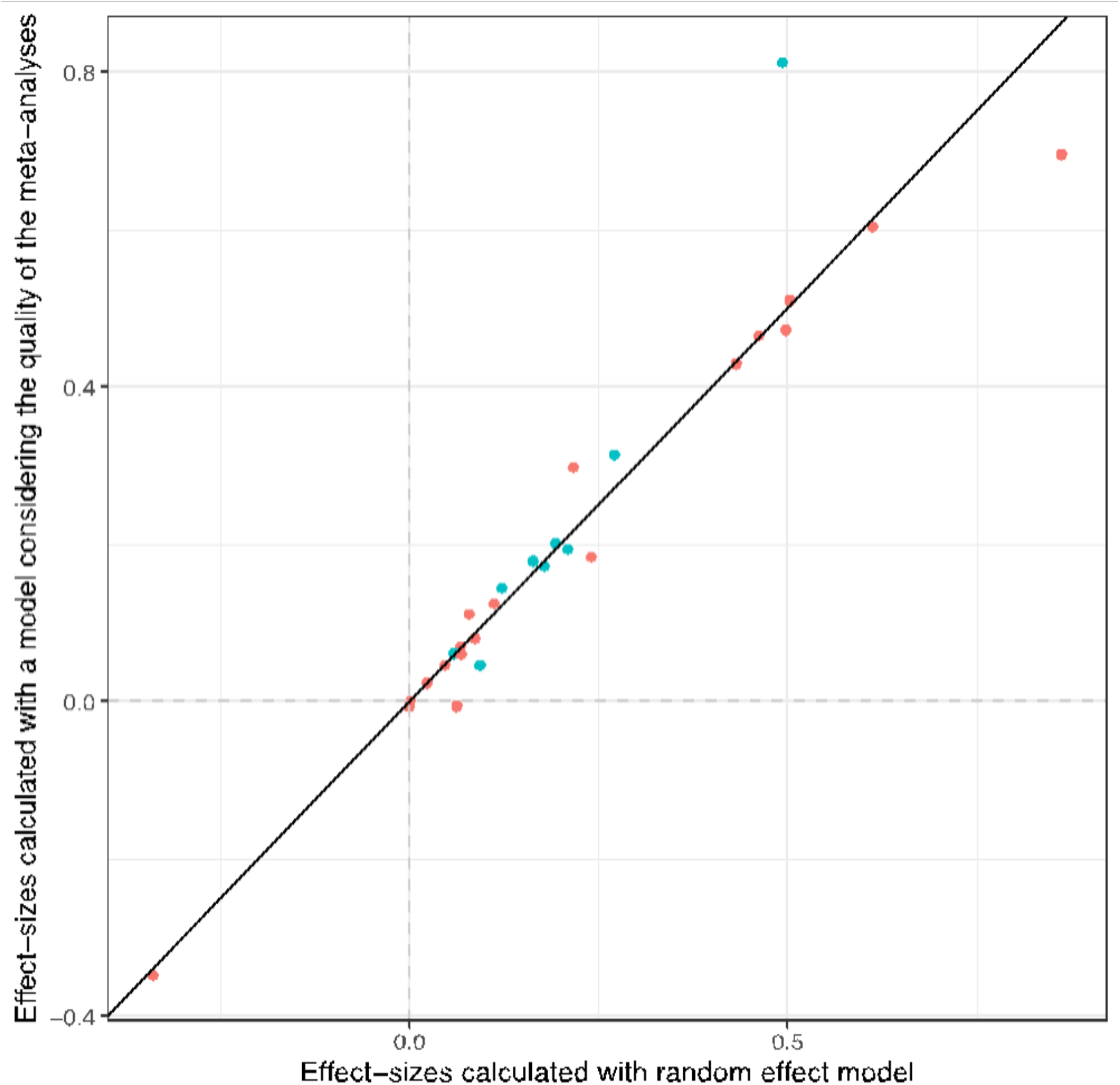
Analysis of the potential impact of the quality of the meta-analyses. The mean effect-sizes calculated with a random effect model (ID and scenario as random effects) are plotted against the same effect-sizes calculated considering the quality of each meta-analyses in the random effect model (see *Doi et al., 2005* for further details). Blue points indicate significant egger tests (p-value < 0.005). Results are presented for the five strategies of crop diversification on the main ecosystem services.

### Text 1

Figure 1

B. Canola fields: Maria Eklind, Svedala, Sweden, under Attribution-ShareAlike 2.0 Generic (CC BY-SA 2.0) licence. Flickr. Wheat field. the aucitron. unkown location. under Attribution-ShareAlike 2.0 Generic (CC BY-SA 2.0) licence. Flickr.

C. Cover crops. USDA NRCS Montana. USA. Public Domain Mark 1.0. flick

D. Agroforestry. Caroline Dangleant, Senegal. CIRAD Copiright.

E. Intercropping. Pierre Sylvie, Paraguay. CIRAD Copiright.

F. Wheat variety mixture. Mathilde Chen. Copiright.

Figure S2

A. Caroline Dangleant. Senegal. CIRAD Copiright.

B. Gmelina arborea, Wikipedia, (CC-BY-SA3.0)

C. Pierre Sylvie. CIRAD Copiright.

D. Unkown location. under Attribution-ShareAlike 2.0 Generic (CC BY-SA 2.0) licence. Flickr.

E. Verena Seufert. Copiright

F. Verena Seufert. Copiright.

**Table S4.**
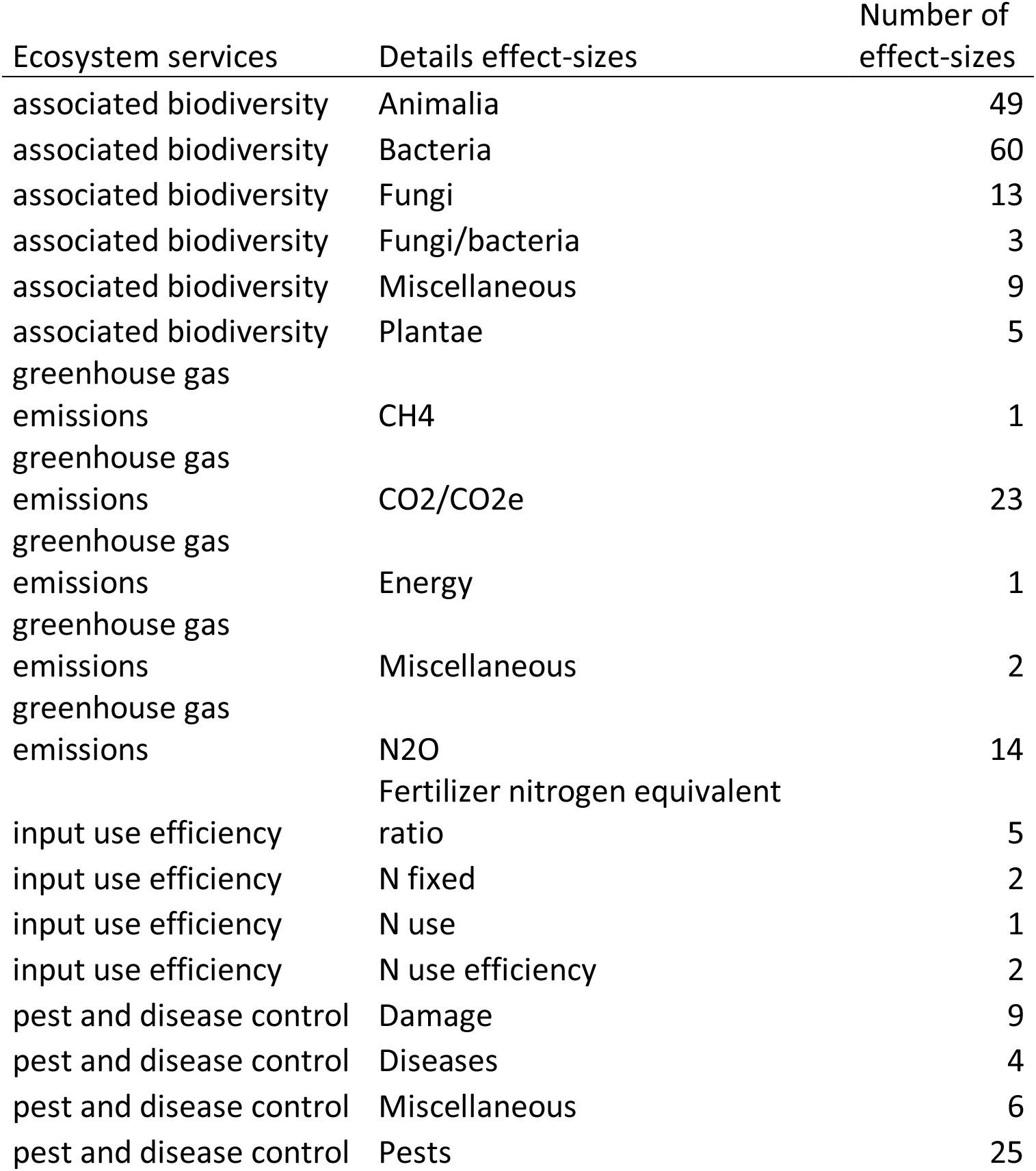
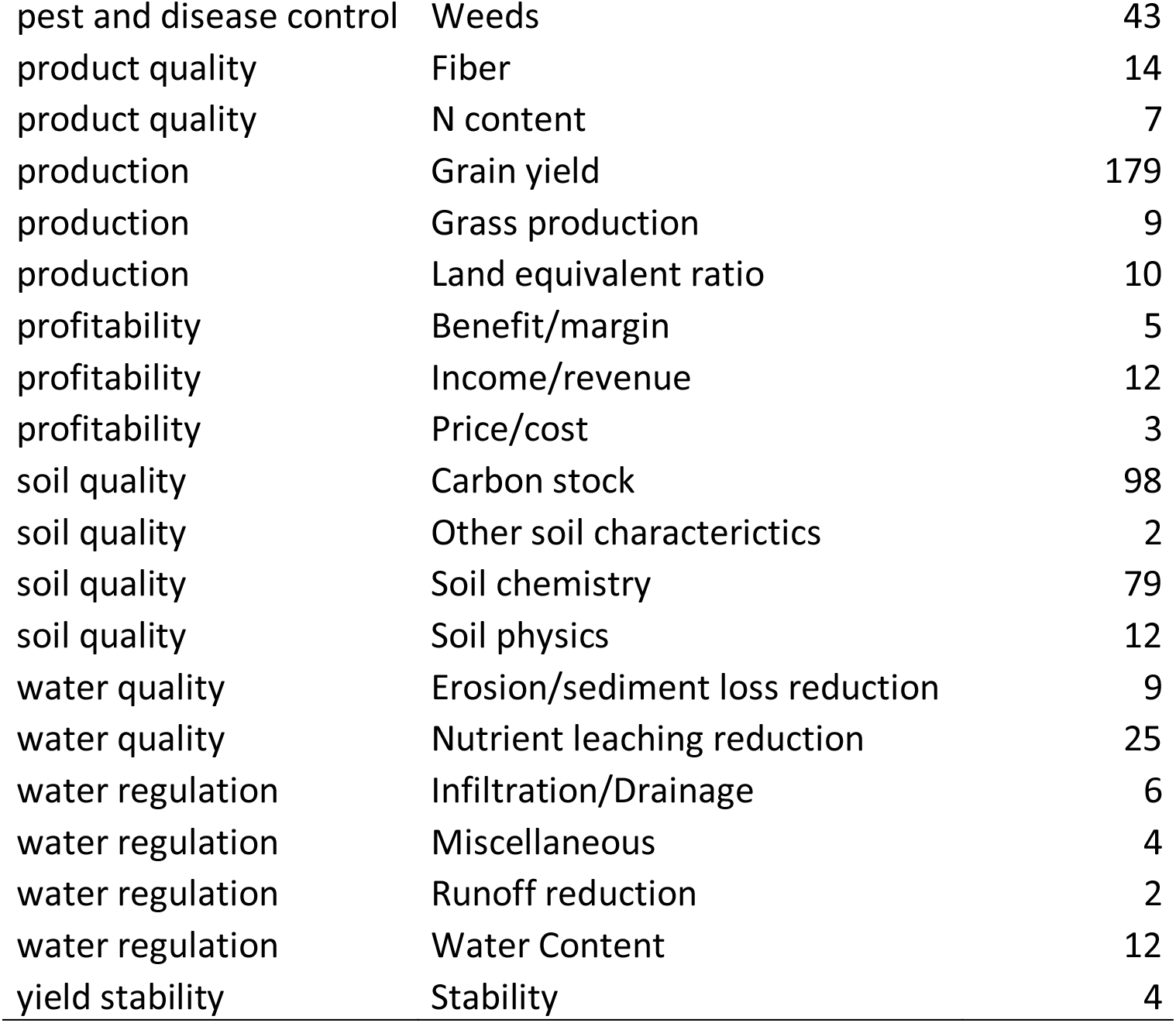

